# Natural Scene Coding Consistency in Genetically-Defined Cell Populations

**DOI:** 10.1101/2025.10.29.685363

**Authors:** Tahereh Toosi, Kenneth D. Miller

## Abstract

Understanding how genetically-defined cell populations encode visual information remains a fundamental challenge in systems neuroscience. While extensive research has characterized individual cell responses to simple stimuli such as static gratings, the population-level coding principles that govern naturalistic visual processing across cell types remain largely unexplored. We analyzed population responses from 43,018 neurons across 12 genetically-defined cell types in 243 mice from the Allen Brain Observatory, comparing representational geometry between natural scenes and static gratings. We found that inhibitory cell populations (VIP, SST, PV) cluster distinctly in representational space almost independent of anatomical location when responding to natural scenes but not static gratings, suggesting preserved cell-type specific computational functions specific for natural scenes. To assess coding capacity of a population of neurons, we developed Inter-Individual Representational Similarity (IIRS), which measures consistency of neural representations across different individuals in response to an ensemble of stimuli. All inhibitory populations showed significantly higher IIRS for natural scenes compared to static gratings, indicating consistent encoding of naturalistic visual features across individuals comparable to excitatory populations (Cux2, Rorb, Rbp4). Parallel analysis of neural networks trained on natural images (ImageNet) with different random initializations revealed similar patterns: models showed higher cross-initialization similarity for naturalistic stimuli compared to static gratings, suggesting that cross-individual consistency emerges when experimental stimuli engage the features that neural circuits are adapted to extract. These findings establish IIRS as a metric for identifying coding capacity in cell populations and reveal that inhibitory cell populations encode consistent aspects of natural scenes across individuals, indicating these circuits may have evolved specialized tuning for naturalistic visual environments.

## 1 Introduction

Research over the last decade has characterized the response properties of different cell types in V1 to simple stimuli like oriented gratings [1–3]. However, the *population-level* coding principles that govern naturalistic visual processing across cell types remain unexplored. Genetic studies have identified cell-type specific vulnerabilities in neurological disorders—parvalbumin (PV) interneuron dysfunction in schizophrenia [4], somatostatin (SST) alterations in Alzheimer’s disease [5], VIP interneurons implicated as a shared vulnerability in autism and schizophrenia [6] and various interneuron abnormalities across psychiatric conditions [7]—yet we cannot predict how these cellular deficits affect visual processing because the computational roles of these cell populations remain poorly understood.

Current approaches to understanding cell-type function focus primarily on single-cell characterizations using artificial stimuli such as drifting gratings and flashing bars [8, 9]. While these studies have revealed important response properties—orientation selectivity, contrast sensitivity, and temporal dynamics—they provide limited insight into how cell populations collectively encode the rich statistical structure of natural visual environments [10]. Moreover, population-level analyses that could reveal distributed coding principles across genetically-defined cell types were less studied, leaving a critical gap between molecular identity and computational function.

We address this limitation by analyzing population-level representations across genetically-defined cell types using large-scale calcium imaging data from the Allen Brain Observatory [3]. Our approach constructs a geometric representation of visual encoding by computing pairwise similarities between population responses from different cell types, cortical layers, and brain regions to both natural scenes and static gratings. We hypothesize that if a cell population encodes meaningful visual features, then the similarity of neural codes between different individual animals should be higher than when response patterns are uninformative. This individual-to-individual representational similarity provides a metric for assessing the coding capacity of genetically-defined populations.

Our analysis reveals three key findings: First, population-level similarity analysis across 100+ distinct celltype combinations (cell-type x depth (cortical layer) x region) shows that inhibitory interneurons cluster distinctly in representational space regardless of anatomical location, suggesting preserved computational functions that transcend local circuit properties. Second, using our individual-to-individual similarity metric, we demonstrate that inhibitory cell populations (SST, VIP, and PV) show significantly higher mouse-to-mouse representational similarity for natural scenes compared to static gratings, indicating consistent encoding of naturalistic visual features. Third, layer-specific analysis of excitatory populations reveals that superficial layer excitatory cells (Cux2 in layers 2/3 and 4, Rorb in layer 4 and Rbp in layer5) also exhibit enhanced similarity for natural scenes. These findings provide a systematic characterization of population-level visual encoding across genetically-defined cell types and establish a framework for linking cellular identity to computational function in naturalistic contexts.

### 1.1 Single-Cell Characterization of Genetically-Defined Cell Types

Extensive research has characterized the response properties of individual genetically-defined neurons in the visual cortex using rigorously-defined but inevitably simplified stimuli. Studies of PV interneurons have revealed their role in providing fast, perisomatic inhibition and generating gamma oscillations [12, 13]. SST interneurons have been shown to provide dendritic inhibition and implement surround suppression [1, 14, 15]. VIP interneurons exhibit disinhibitory effects through their preferential targeting of other interneuron populations [16, 17]. Excitatory cell types have been characterized by their laminar distribution and projection targets, with Cux2-expressing neurons in superficial layers showing distinct response properties from deeper layer Nr5a1-expressing cells [18]. While these studies have established fundamental response properties and circuit connectivity, they focus primarily on single-cell characterizations using artificial stimuli such as drifting gratings and oriented bars, leaving population-level coding of naturalistic scenes largely unexplored.

### 1.2 Population-Level Neural Coding Across Cell Types

Population-level analyses have revealed distributed coding principles that cannot be observed at the single-cell level. Studies using dimensionality reduction techniques have shown that neural populations encode stimulus information in low-dimensional manifolds [19]. Representational similarity analysis (RSA) has been used to compare neural representations across brain areas and species [20, 21]. Recent work has begun to examine population responses across cell types, with studies showing that inhibitory and excitatory populations exhibit distinct temporal dynamics and stimulus selectivity [1, 22]. However, these approaches typically aggregate responses within broad cell classes (excitatory vs. inhibitory) rather than examining how specific genetically-defined populations contribute to the distributed coding of naturalistic stimuli.

### 1.3 Cross-Subject Neural Analysis and Alignment

The challenge of comparing neural representations across different subjects has received increasing attention. Thobani et al. [23] introduced inter-animal transforms as a framework for model-brain alignment, focusing on finding optimal mappings between corresponding anatomically defined brain areas across subjects. Their approach identifies transform classes that map responses between subjects within the same brain area, assuming that such mappings should exist and optimizing alignment quality. Other work has addressed cross-subject variability in neural population structure using canonical correlation analysis and other linear alignment methods [23, 24].

Our work takes a fundamentally different approach to cross-subject analysis. Rather than assuming that cross-subject similarity should exist for anatomically-defined regions, discarding the significant individual variability across regions, we use the presence or absence of cross-subject similarity as a diagnostic tool for coding capacity in (genetically-defined) cell populations. We compare inter-individual representational similarity between stimulus distributions where we expect meaningful coding (natural scenes) versus distributions where we expect limited coding (artificial gratings). This approach remains agnostic to anatomical boundaries, recognizing that individual variation in brain structure [18] means that functional populations may not align perfectly across subjects based on anatomical location alone. By using cross-subject consistency as an emergent property rather than an optimization target, we can assess whether genetically-defined cell populations exhibit selective encoding of specific stimulus statistics.

## 2 Results

### 2.1 Population-Level Similarity Analysis Reveals Distinct Cell-Type Organization

To understand how genetically-defined cell populations collectively encode visual information, we computed pairwise similarities between population responses across all combinations of cell type, cortical layer, and brain region. Using Centered Kernel Alignment (CKA) to measure representational similarity, we analyzed over 100 distinct population combinations (cre-line x cortical layer x region) responding to natural scenes (Figure 2A). The resulting similarity matrix reveals a striking organizational principle: inhibitory cell populations cluster together regardless of their anatomical location (Figure 2A). This clustering pattern suggests that cell-type identity may be more fundamental to computational function than regional boundaries. Hierarchical clustering analysis confirmed that inhibitory populations (VIP, SST, and PV) form distinct groups that are clearly separated from excitatory populations.

**Figure 1:**
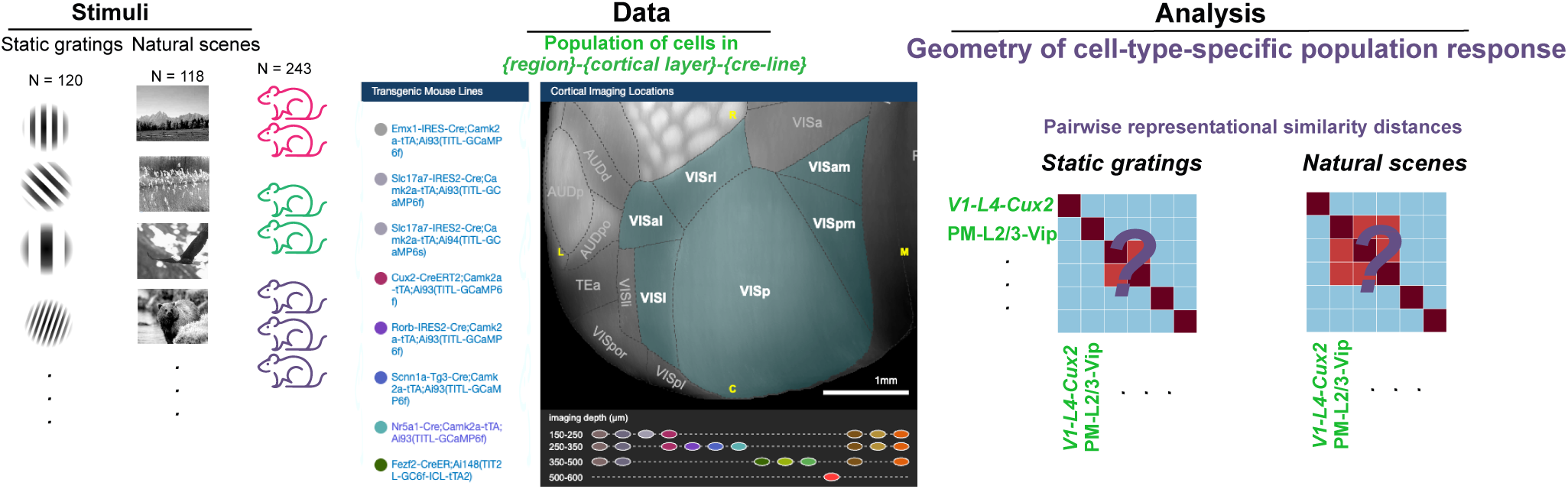
Cell-type specific population codes for natural scenes. Systematic investigation of cell-type specific visual processing across two data distributions: natural scenes (n=118) and static gratings (systematically-varied orientation and spatial frequency, n=120). **Stimuli:** Stimulus sets including static gratings and natural scenes are shown to mice during neural recordings. **Data:** Neural responses are recorded from genetically-identified cell populations across 4 cortical layers and 6 visual areas. Brain map adapted from Allen Brain Observatory Visual Coding dataset [11] (Image credit: Allen Institute for Brain Science, https://observatory.brain-map.org/visualcoding/). **Analysis:** We constructed similarity matrices of cell populations to quantify representational geometry to reveal which cell types respond similarly to the same visual inputs, essentially mapping which neural populations “see” the world in comparable ways and identifying functional relationships across genetically-defined cell types.

**Figure 2:**
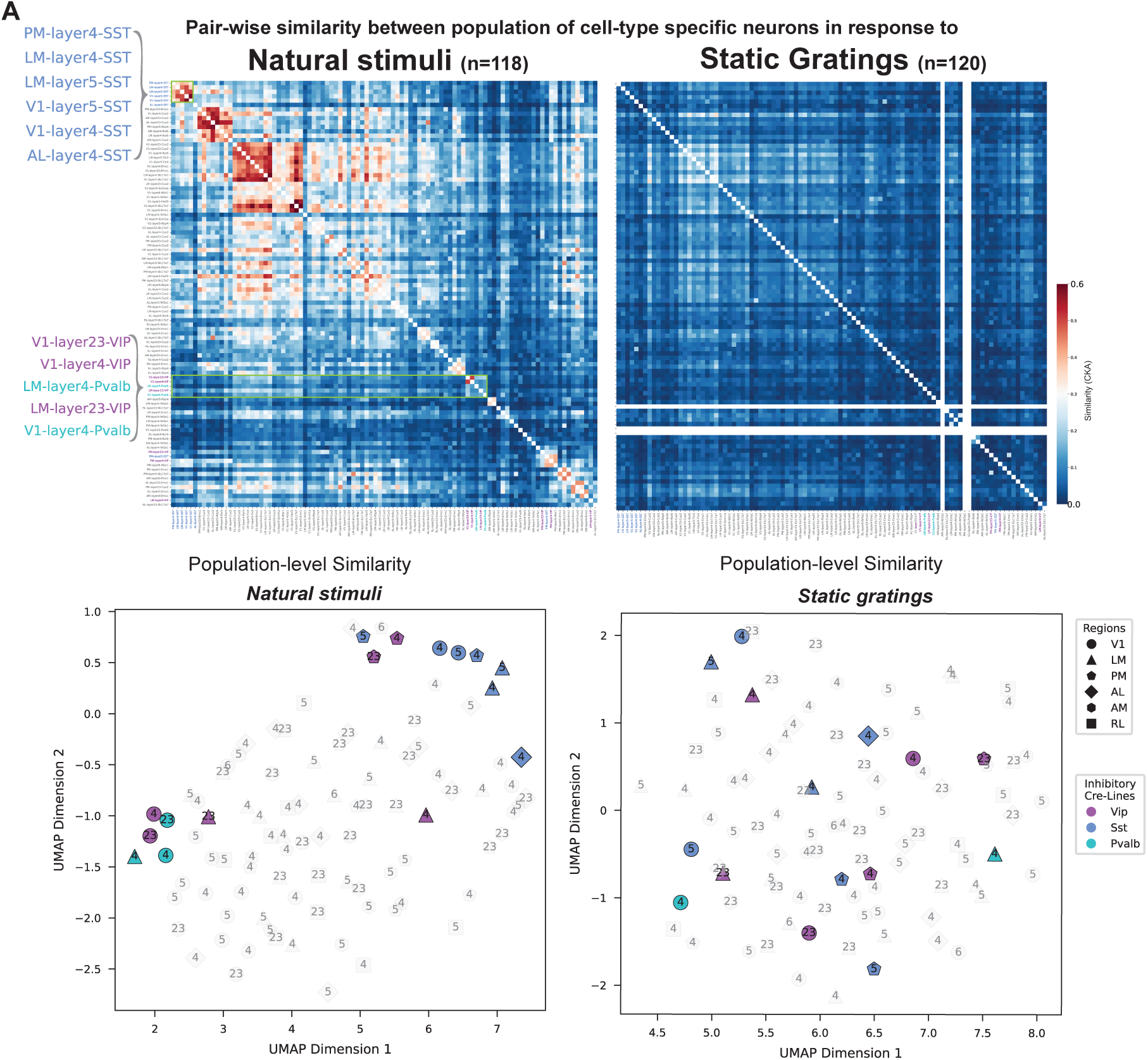
A Population-level similarity across region. *×* **layer** *×* **cre-line** (e.g. PM-layer4-SST) combinations for 118 natural scenes (left) and 120 static gratings (right). Metric: linear CKA. Both panels use identical row/column ordering based on hierarchical clustering of natural scene responses. Diagonal (self-similarity) entries removed for visualization. Blank entries indicate populations absent from static grating dataset (n=3). Color-coded labels indicate cre-line identity. **B Left:**UMAP visualization of population responses to static gratings **Right:** natural scenes. Numbers indicate cortical layers (23: layers 2/3, 4-6: respective layers). Colored markers represent inhibitory neuron populations (cyan: PV, blue: SST, purple: VIP), while gray markers show excitatory populations. Marker shapes indicate different visual areas (circles: V1, triangles: LM, pentagons: PM, diamonds: AL, etc., side panels show magnified views of regions outlined in green boxes.). The pattern of clustering is robust to different UMAP runs (Figure S3). Only cells with split-half reliability*>*0.4 were included for both types of stimuli to ensure fair comparison. Same neurons were recorded under both conditions and data for each population was pooled across available cre-line mice [3] .

To visualize these high-dimensional relationships, we applied UMAP dimensionality reduction to the population similarity data. The resulting embedding shows that inhibitory cell types maintain their distinct clustering across both natural scenes and static gratings (Figure 2B). Notably, while the overall organization is preserved, the relative positions of populations shift between stimulus types, indicating stimulus-dependent reorganization of representational relationships. The consistency of inhibitory clustering across multiple UMAP initializations confirms the robustness of this organizational principle (Supplementary Figure S3).

### 2.2 Inter-Individual Representational Similarity Reveals Stimulus-Selective Coding

To assess whether the observed population structure reflects meaningful coding rather than general responsiveness, we developed Inter-Individual Representational Similarity (IIRS), a metric that quantifies the consistency of neural representations across different individual mice. We hypothesized that if a cell population encodes specific stimulus features, then the similarity of neural codes between different mice should be higher for stimuli that engage the population’s specialized function.

o assess stimulus-selective coding, we computed IIRS distributions for each cell type under both stimulus conditions and compared them using Wilcoxon rank-sum tests (Figure 3). Each IIRS distribution comprised similarity values from all pairwise comparisons between mice within a cell type (excluding self-comparisons). Sample sizes were: SST (n=27 mice, 351 pairs), VIP (n=24 mice, 276 pairs), PV (n=8 mice, 28 pairs), Cux2 (n=38 mice, 703 pairs), Rbp4 (n=23 mice, 253 pairs), and Rorb (n=24 mice, 276 pairs) (Supplementary FigureS4). All p-values were Bonferroni-corrected for multiple comparisons across cell types (*α* = 0.05/6 = 0.008). All three inhibitory populations showed significantly higher inter-individual similarity for natural scenes compared to static gratings (SST: W = 2468.0, p *<* 0.001; VIP: W = 2208.0, p *<* 0.001; PV: W = 1427.0, p *<* 0.001). Excitatory populations showed similar patterns. Superficial and mid-layer excitatory cells all exhibited significantly higher IIRS for natural scenes (Cux2: W = 3486.0, p *<* 0.001; Rorb: W = 2208.0, p *<* 0.001; Rbp4: W = 1722.0, p *<* 0.001). In contrast, Nr5a1-expressing neurons (Supplementary Figure S5) showed no significant difference in IIRS between natural scenes and static gratings (W = 1076.0, p = 0.337), consistent with their specialization for dynamic rather than static visual features.

**Figure 3:**
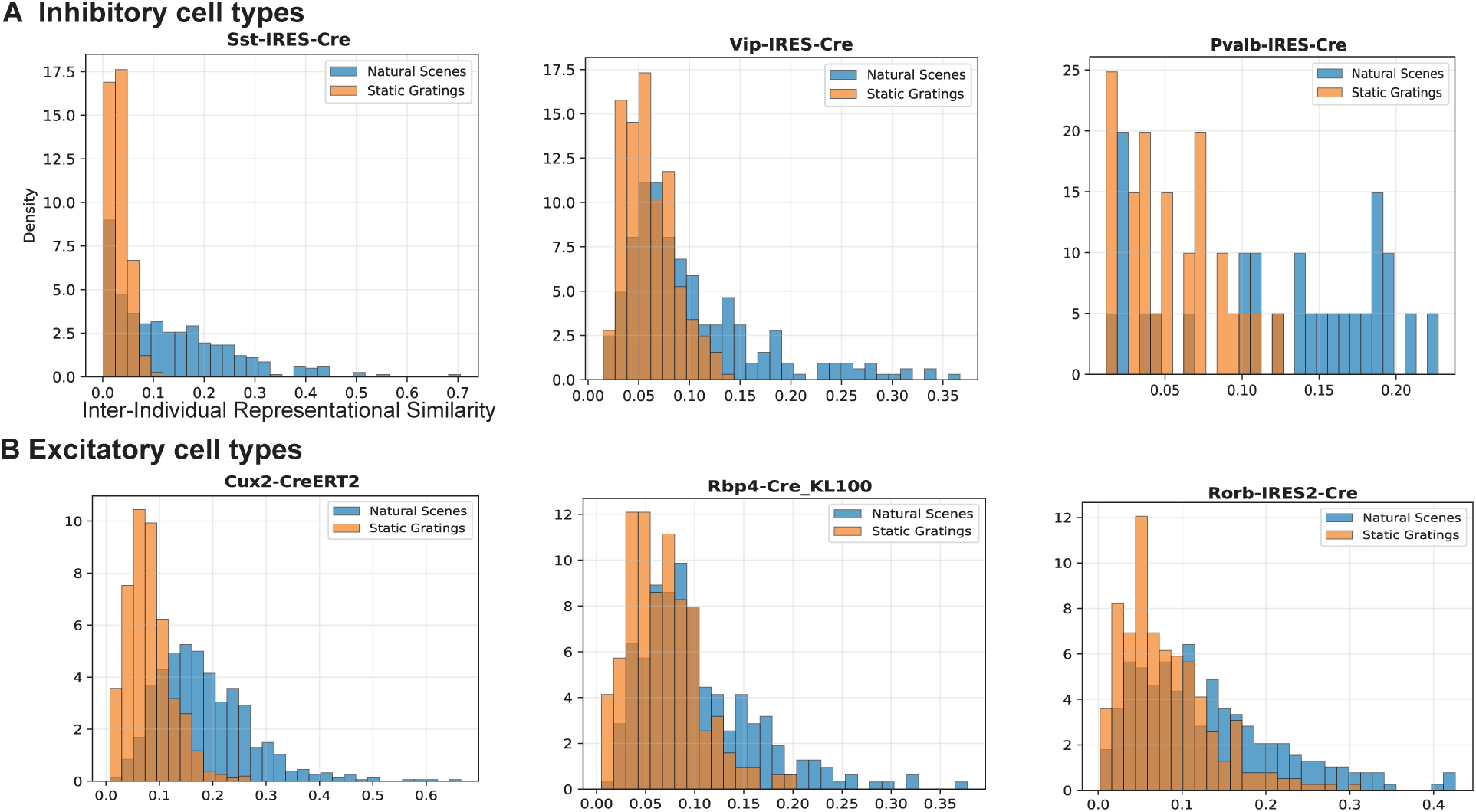
Inter-individual representational similarity reveals stimulus-selective coding capacity across genetically-defined cell types. Distribution of CKA similarity values between individual mice for natural scenes (blue) and static gratings (orange). **Top row:** Inhibitory populations (SST, VIP, PV) show consistently higher inter-individual similarity for natural scenes, indicating robust encoding of naturalistic features. **Bottom row:** Excitatory populations show similar effect with varing degree. Each distribution represents similarity values from all pairwise comparisons between mice (excluding selfcomparisons). For the original similarity matrices, see Supplementary Figure S4

The IIRS distributions reveal that natural scenes engage coding mechanisms in specific cell populations that are not activated by artificial gratings, providing direct evidence for stimulus-selective population coding in genetically-defined cell types.

### 2.3 Computational Validation: Individual Differences as Initialization Variation

To test whether stimulus-selective representational consistency reflects general principles of learning rather than biological-specific mechanisms, we examined neural networks with different random initializations as a computational model of individual variation. This framework is motivated by fundamental parallels between biological and artificial systems: individual mice share the same genetic “architecture” and visual experience but develop slightly different neural organizations due to stochastic developmental processes, much like neural networks with identical architectures and training that develop different representations due to random weight initialization. This choice of modeling recently was used to explain individual differences in humans [25].

We compared two ResNet50 models trained identically on ImageNet for 1000-way object classification but with different initializations, testing them on the same natural scenes and static gratings used in our biological analyses. Consistent with our biological findings, networks showed higher cross-initialization similarity for natural scenes compared to static gratings (Figure 4), albeit with different distributions. This result suggests that stimulus-selective representational consistency emerges from the degree of match between training and test distributions: networks converge to more similar solutions when processing stimuli that share statistical structure with their training data.

**Figure 4:**
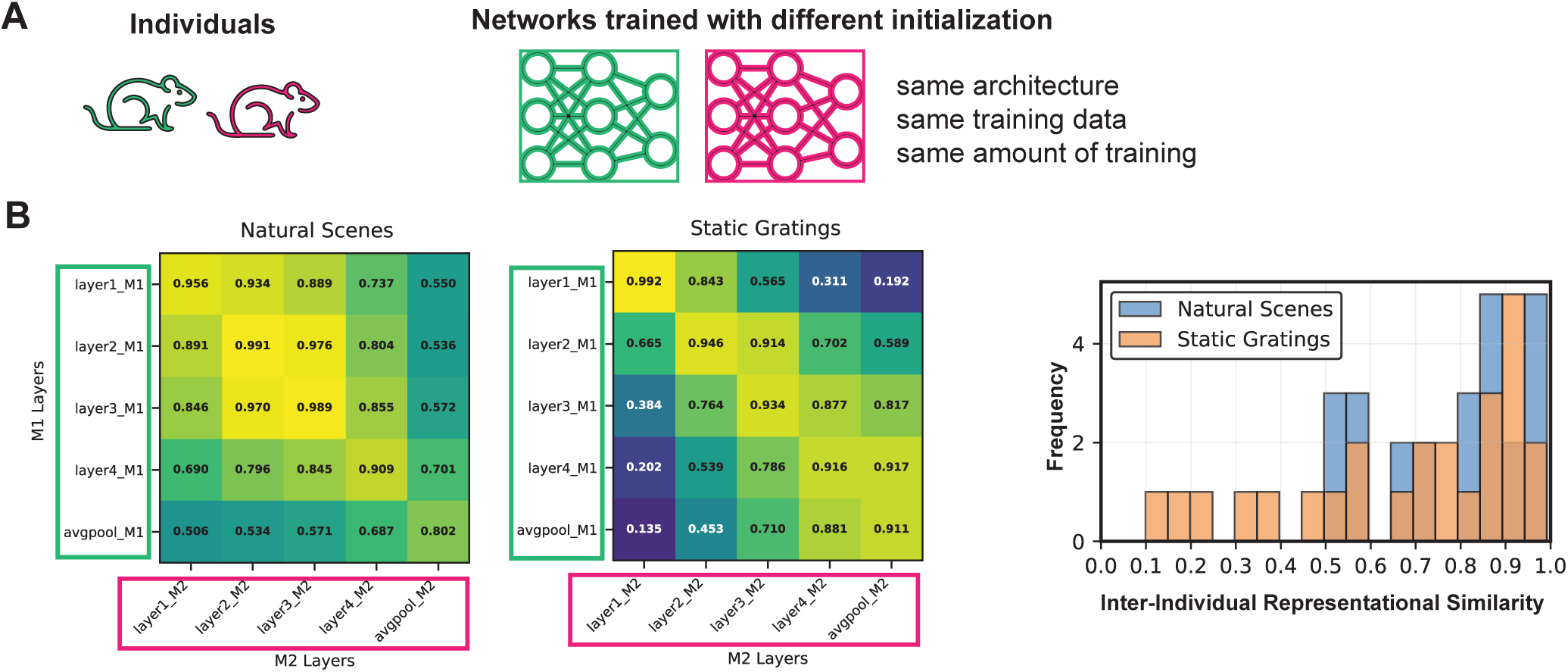
Individual animals can be modeled as instances of a model class with different initializations. **A.** Conceptual framework: Individual biological organisms (left) can be understood as instances of the same “model class” (shared genetic architecture and developmental programs) with different random initializations (genetic variation, environmental factors). Similarly, neural networks (right) with identical architecture trained on the same dataset (ImageNet) can achieve comparable performance while developing different internal representations due to different weight initializations. **B.** Cross-model CKA similarity matrices comparing two ResNet50 models trained with different initializations, tested on static gratings (left) and natural scenes (middle), the same stimuli used in neural analyses from [3]. Both networks were trained on ImageNet natural images.

This parallel between biological individual differences and computational initialization differences supports the interpretation that our biological findings reflect fundamental properties of learning systems rather than mouse-specific neural mechanisms. Complete methodological details are provided in Supplementary Methods S4.

## 3 Discussion

This study establishes Inter-Individual Representational Similarity (IIRS) as a novel diagnostic tool for identifying coding capacity in genetically-defined neural populations. Our analysis of over 43,000 neurons across multiple cell types in mouse visual cortex reveals that stimulus-selective representational consistency represents a fundamental organizing principle that distinguishes meaningful coding from general responsiveness.

The finding that inhibitory populations show robust cross-individual consistency for natural scenes but not artificial gratings extends traditional views by suggesting inhibitory populations contribute more actively to naturalistic coding than previously appreciated. The parallel patterns observed in excitatory populations indicate that this organizational principle extends across cell types, with layer-specific variations reflecting distinct computational specializations. The consistent differences between population-level responses to natural scenes versus static gratings suggests that conclusions based on artificial stimulus processing may not generalize to naturalistic visual environments. This limitation is verified in our neural network analysis, where models trained on natural images showed clear representational differences when tested on natural scenes compared to static gratings. These findings highlight a critical gap in traditional neuroscience approaches: insights derived from simplified, parametric stimuli may inadequately capture the computational principles governing real-world sensory processing, necessitating a shift toward more naturalistic experimental paradigms.

The computational validation using neural networks with different initializations suggests these patterns may reflect general properties of learning systems, not specific to this dataset. Networks show higher representational consistency when processing stimuli that match their training distribution, paralleling the biological observation that populations show enhanced consistency for naturalistic inputs.

These findings have broader implications for understanding neural population codes and individual variation in brain function. The IIRS framework can be applied to other datasets and brain regions to identify coding populations and assess the functional significance of genetic or experiential differences. By moving beyond single-cell characterizations to population-level representational geometry, this approach provides a bridge between molecular identity and computational function that will advance our understanding of how diverse cell types contribute to sensory processing.

## 4 Methods and Materials

We analyzed neural activity data from the Allen Brain Observatory Visual Coding 2P dataset [3], which contains two-photon calcium imaging recordings from genetically-defined cell populations in mouse visual cortex. We constructed a comprehensive similarity matrix across all cre-line-cortical-layer-region population combinations by computing pairwise Centered Kernel Alignment (CKA)[26] between population responses. This analysis revealed the geometric organization of visual representations across genetically-defined cell types. Then using UMAP [27] dimensionality reduction (for visualization purposes only, all scientific conclusions were drawn from the original distance matrix). Full details of the population-level analysis pipeline are provided in Supplementary Methods S2.

### Inter-Individual Representational Similarity (IIRS)

To assess the coding capacity of genetically-defined populations, we developed a metric that compares representational consistency across individual mice. For each cell type, we computed CKA similarity between population responses from different mice, comparing natural scenes versus static gratings to identify stimulus-selective encoding. This contrastive approach uses cross-subject consistency as a diagnostic for meaningful coding rather than stimulusunspecific selectivity. IIRS was computed as:

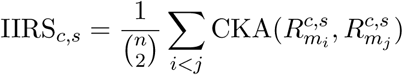

where 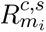 represents the response matrix for mouse *i*, cell type *c*, and stimulus set *s*. Complete IIRS methodology and statistical analysis procedures are detailed in Supplementary Methods S3.

## A Supplementary Material

### A.1 S1. Dataset and Preprocessing

#### A.1.1 Allen Brain Observatory Visual Coding 2P Dataset

We analyzed data from the Allen Brain Observatory Visual Coding 2P dataset [3], which contains twophoton calcium imaging recordings from mouse visual cortex during presentation of standardized visual stimuli. The dataset includes recordings from six visual areas (VISp, VISl, VISal, VISpm, VISam, VISrl) across multiple cortical layers, with cells genetically labeled using cre-driver lines. Data were accessed through the Allen SDK (https://allensdk.readthedocs.io/) and the Brain Observatory Cache system.

##### Stimulus Sets

We analyzed responses to two stimulus categories that represent different statistical distributions:

##### Natural scenes

118 natural images from the Allen Brain Observatory dataset, presented for 250ms each with approximately 50 trials per image. Images were grayscale, contrast-normalized, and spanning 120° × 95° of visual space. Trials were randomized, with blank sweeps appearing approximately once every 100 images. These stimuli contain rich, naturalistic statistical structure including 1/f spatial frequency distributions and complex spatial correlations.

##### Static gratings

120 sinusoidal gratings systematically varying across three dimensions:

- Orientation: 6 values (0°, 30°, 60°, 90°, 120°, 150°)
- Spatial frequency: 5 values (0.02, 0.04, 0.08, 0.16, 0.32 cycles/degree)
- Phase: 4 values (0, 0.25, 0.5, 0.75)

Each grating was presented for 250ms with approximately 50 trials per condition at 80% contrast. Trials were randomized, with blank sweeps presented approximately once every 25 trials. These stimuli represent artificial, parametrically controlled visual patterns with sparse statistical structure.

##### Quality Control Criteria

Strict quality control was applied to ensure reliable population analyses:

- **Minimum cell count**: 10 cells per mouse for each analyzed cre-line-region-depth combination
- **Response reliability**: Trial-averaged responses computed only for cells with valid recordings across all stimulus presentations
- **Session quality**: Excluded sessions flagged with technical artifacts based on Allen Institute quality metrics
- **Cortical area coverage**: Included only cell types with representation across multiple visual areas in IIRS analysis

#### A.1.2 Population Definition and Grouping

We defined distinct neural populations using the combination *{*cre-line*}*-*{*cortical-layer*}*-*{*region*}*:

##### Cre-line

Neurons were classified by cell type using transgenic Cre-driver lines that label genetically defined populations. The study included 12 distinct cell populations comprising both excitatory and inhibitory neurons. Excitatory populations included: Cux2-CreERT2/wt (superficial layers), Rorb-IRES2-Cre/wt (layer 4), Scnn1a-Tg3-Cre/wt (layer 4), Rbp4-Cre KL100/wt (layer 5), Nr5a1-Cre/wt (layer 4), Tlx3-Cre PL56/wt (layer 5), Ntsr1-Cre GN220/wt (layer 6), Slc17a7-IRES2-Cre/wt (broad excitatory), Fezf2-CreER/wt (layer 5), and Emx1-IRES-Cre/wt (broad excitatory). Inhibitory populations included: VIP-IRES-Cre/wt (VIP+ interneurons), SST-IRES-Cre/wt (somatostatin+ interneurons), and PV-IRES-Cre/wt (parvalbumin+ interneurons). The cell counts across mice for each population is provided in Figure S1.

##### Cortical layer

Inferred from imaging depth using Allen Institute conventions:

- Layer 2/3: 175m depth
- Layer 4: 230-300m depth
- Layer 5: 350-375m depth
- Layer 6: 500m depth

##### Region

Six visual areas (VISp/V1, VISl/LM, VISal/AL, VISpm/PM, VISam/AM, VISrl/RL)

This yielded over 100 distinct population combinations, with variable representation across combinations based on natural cre-line distributions.

#### A.1.3 Centered Kernel Alignment (CKA)

We computed population similarities using linear CKA, which measures representational similarity while being invariant to orthogonal transformations:

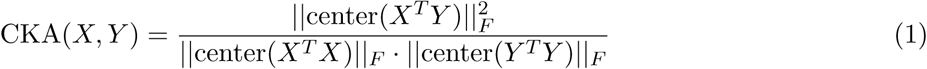

where:

- *X, Y* are neural response matrices (n stimuli × n neurons)
- center(*·*) applies row and column centering: center(*M*) = *HMH* where 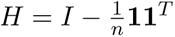
- *|| · ||_F_* denotes the Frobenius norm

CKA values range from 0 (no similarity) to 1 (perfect similarity after orthogonal transformation).

### A.2 Representational Similarity Analysis

Population-level similarities were quantified using Centered Kernel Alignment (CKA) with linear kernels [26], which measures the similarity between representational geometries while being invariant to orthogonal transformations. For each pair of populations, CKA similarity was computed between their stimulus response vectors. The resulting similarity matrices were visualized using Uniform Manifold Approximation and Projection (UMAP) with precomputed distance matrices (1 - CKA similarity) to reveal organizational principles in high-dimensional representational space.

### A.3 UMAP Analysis

To visualize the high-dimensional representational similarities between cell-type populations, we employed Uniform Manifold Approximation and Projection (UMAP) [27], a nonlinear dimensionality reduction technique that preserves both local and global structure in high-dimensional data. UMAP was applied to the precomputed distance matrices derived from CKA similarities (distance = 1 - CKA similarity) to project the population relationships into a two-dimensional visualization space. UMAP projections were computed using standard parameters: 15 nearest neighbors, minimum distance of 0.1, and precomputed metric applied to the precomputed distance matrices. These parameters were selected to balance preservation of local neighborhood structure while maintaining global organizational patterns. To ensure that the observed organizational patterns were not artifacts of UMAP’s stochastic optimization procedure, we systematically tested robustness across multiple random initializations (Figure S3). UMAP embeddings were computed for 4 different random seeds for both natural scenes and static gratings stimulus conditions.

### A.4 Deep neural network modeling

To investigate whether the stimulus-dependent differences observed in biological neural populations might emerge from general principles of visual representation learning, we modeled each individual mouse using identical ResNet50 architectures [28]. In this framework, each genetically defined cell-type population (e.g., VISp-layer4-Cux2, VISpm-layer5-SST) was conceptualized as corresponding to a distinct layer within the network architecture. Although the network layers are not exclusively inhibitory or excitatory like their biological counterparts, this abstraction allows us to examine whether similar stimulus-dependent organizational principles emerge from computational constraints alone.

All ResNet50 models were trained on ImageNet [29] for image classification using identical training procedures and hyperparameters. Crucially, each model instance was exposed to exactly the same amount of training data, differing only in random weight initialization, thereby controlling for experience-dependent factors while allowing initialization-dependent representational differences to emerge. This design enables direct comparison with the biological data, where individual mice share the same genetic architecture and visual experience but may develop slightly different functional organizations.

Model checkpoints were obtained from the Hugging Face model repository [30], providing multiple independently trained ResNet50 instances with different random initializations. Feature representations were extracted from intermediate layers and compared using the same CKA-based similarity analysis applied to the biological neural data, allowing direct quantitative comparison between artificial and biological visual processing systems.

**Figure S1:**
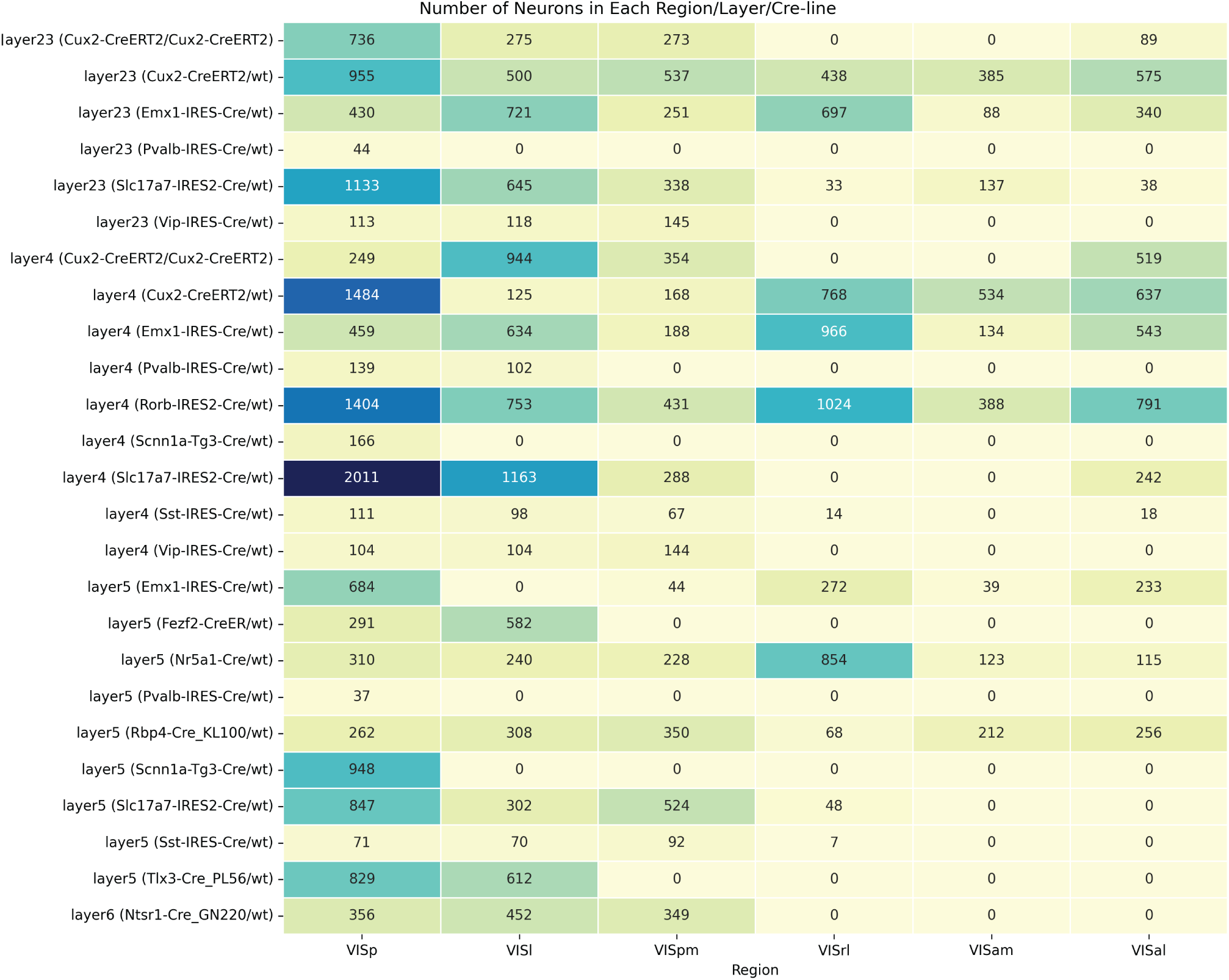
Cell count distribution across genetically-defined populations, cortical layers, and visual areas. Heatmap showing the number of neurons for each combination of cell type (rows), cortical layer, and visual area (columns) included in the analysis after quality control. Cell counts represent the total number of neurons across all mice for each population combination.

## B Supplementary result figures

### B.1 Similarity matrix for Static gratings

**Figure S2:**
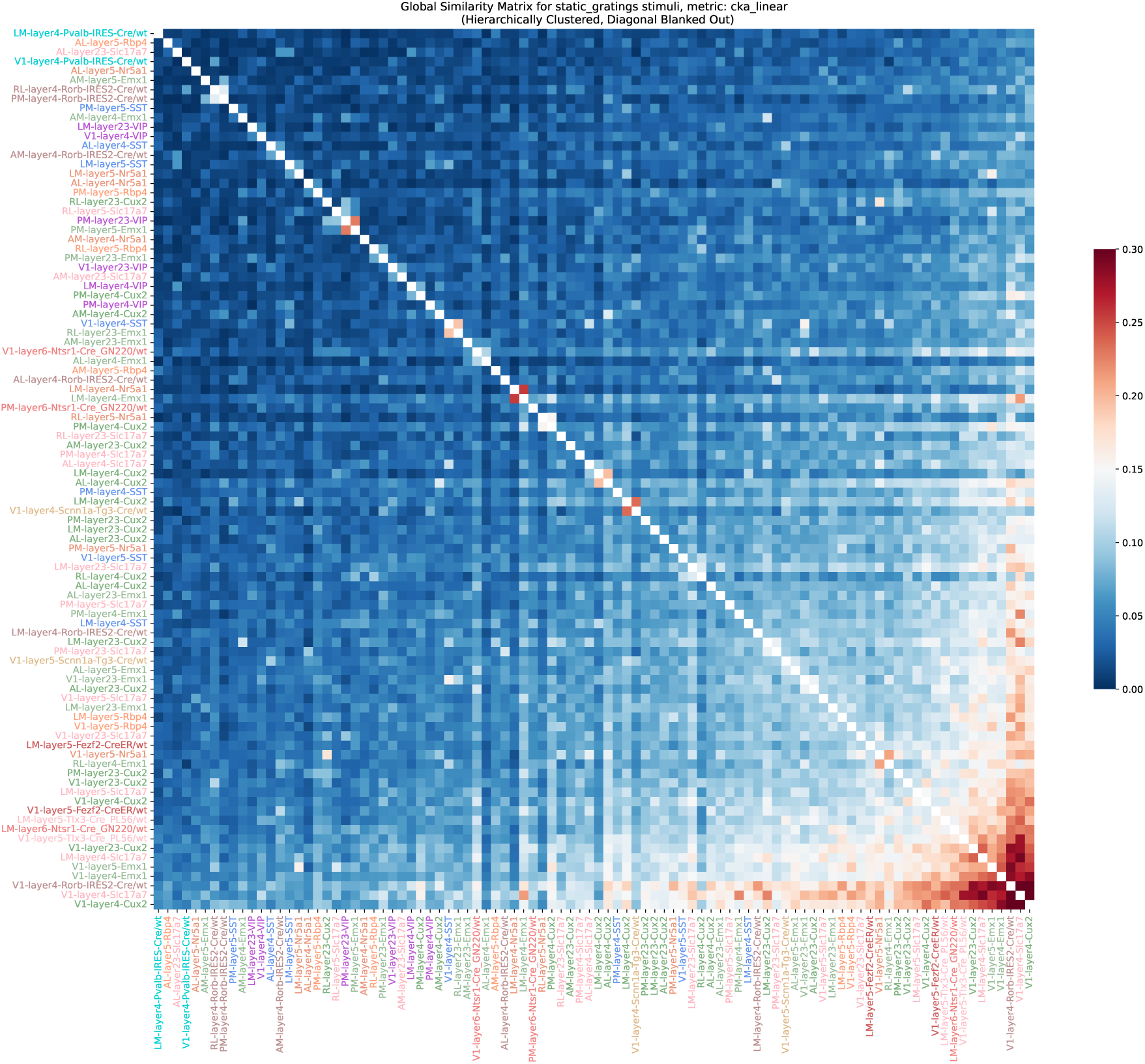
Population-level similarity analysis across combination of region x cortical-layer x cre-line (e.g. V1-layer4-PV) for pairwise population responses to static gratings (metric: linear CKA, 120 systematically varying static grating images, the order was based on hierarchical clustering of the same data)

### B.2 Accounting for UMAP stochastic optimization

**Figure S3:**
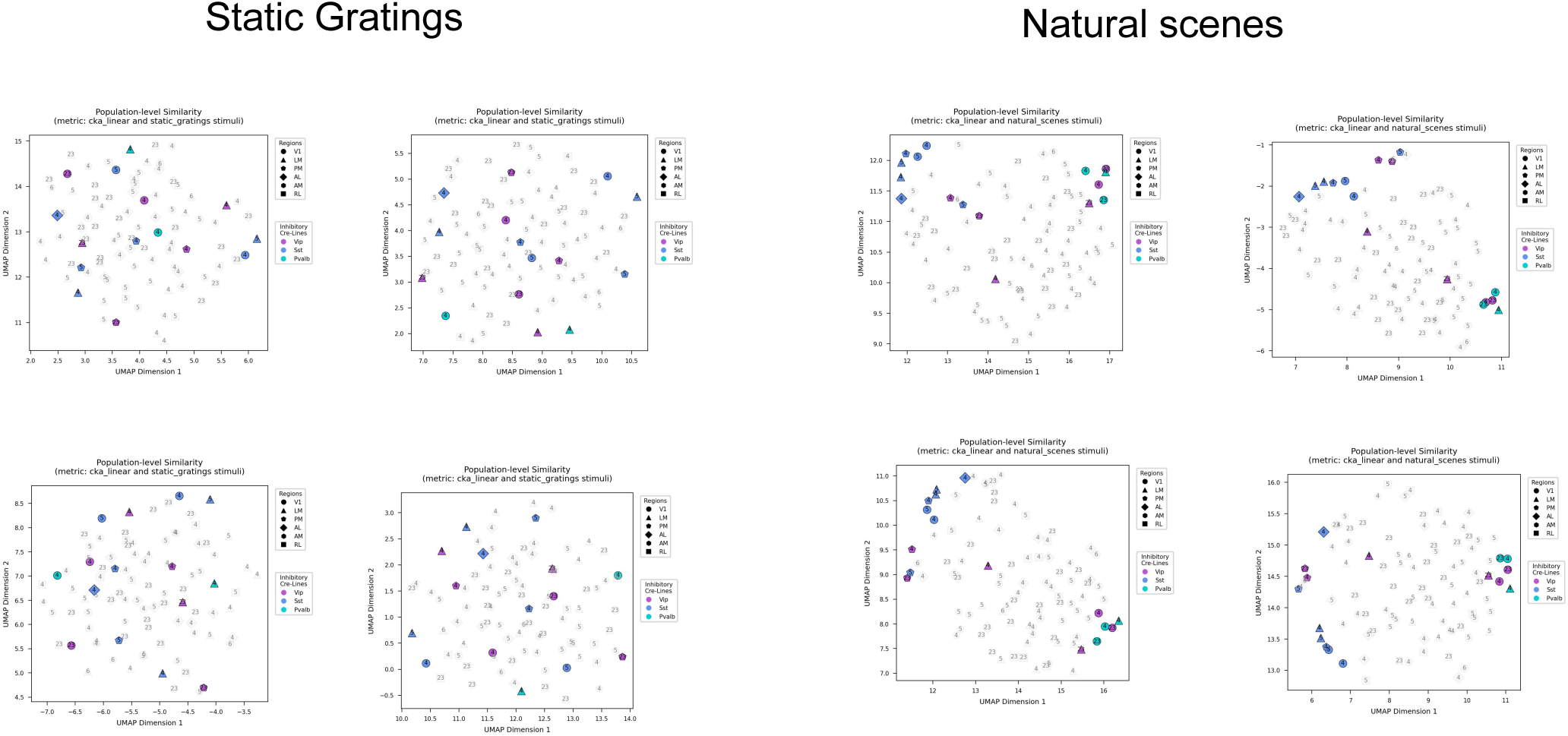
Robustness of UMAP embedding across multiple random initializations. UMAP visualizations of population-level similarities computed with four different random seeds for static gratings (left) and natural scenes (right). Each panel shows the same data projected with a different random initialization of the UMAP algorithm (seeds 1-4). Despite stochastic optimization differences, key organizational principles remain consistent across runs: inhibitory populations (VIP, SST, PV; colored markers) maintain distinct clustering in natural scenes but show more dispersed organization for static gratings. We only used UMAP (and tSNE, results not shown) as visualization tools and the similarity of inhibitory neurons can be seen in original distance matrix. Gray markers represent excitatory populations; shapes indicate visual areas (circles: V1, triangles: LM, diamonds: AL, pentagons: PM, etc.); numbers indicate cortical layers.

### B.3 Inter-individual similarity matrices for each cre-line

**Figure S4:**
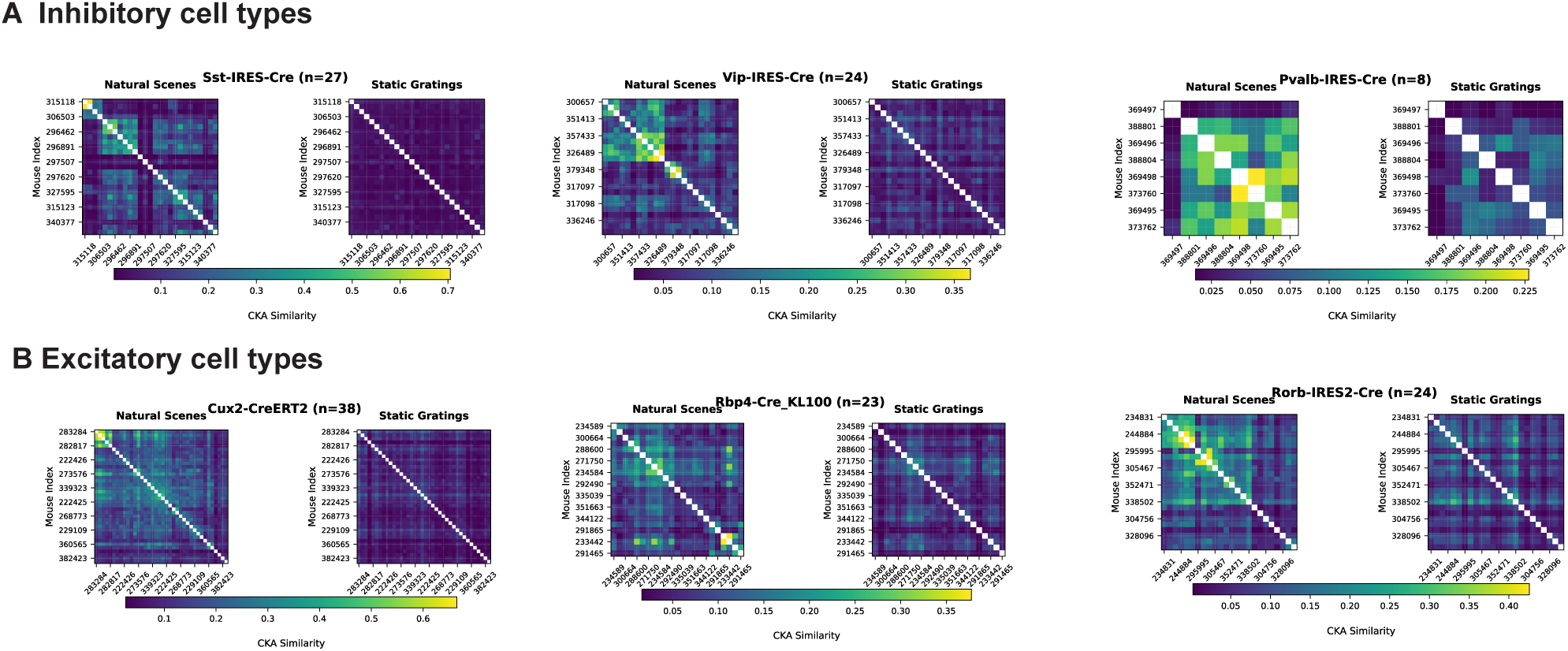
Complete inter-individual similarity matrices and distributions for inhibitory and excitatory cell types. **A.** Pairwise CKA similarity matrices between individual mice for inhibitory populations (SST, VIP, PV) showing natural scenes (left) vs. static gratings (right) for each cell type. Each matrix element represents similarity between two individual mice, with sample sizes indicated (n = number of mice). Warmer colors indicate higher similarity. **B.** Corresponding matrices for excitatory populations (Cux2, Rbp4, Rorb). Individual mouse similarity matrices reveal the underlying structure captured by the IIRS metric, with natural scenes consistently producing higher cross-individual similarities compared to static gratings across both inhibitory and excitatory populations. The complete distributions shown here provide the raw data from which the upper triangular values were extracted for the histogram analysis presented in Figure 3. Diagonal elements (self-similarity = 1.0) are excluded from IIRS calculations.

#### B.3.1 Direction-Selective Populations Show Different Coding Patterns

Nr5a1-expressing neurons showed no significant difference in IIRS between natural scenes and static gratings ((W = 15273, p = 0.4964); Figure S5). This pattern differs from other excitatory populations and likely reflects the specialized response properties of Nr5a1 cells. These neurons are known for their strong direction selectivity and preferential responses to dynamic visual features rather than static spatial patterns [31].

The lack of enhanced natural scene coding in Nr5a1 populations may indicate that their computational specialization lies in processing temporal dynamics—information that is absent in both our static natural scenes and static gratings. This suggests that different cell populations may be specialized for different aspects of visual processing: spatial pattern encoding (as seen in Cux2, Rorb, Rbp4) versus motion processing (Nr5a1), with our static stimulus paradigm primarily engaging the former. We plan to investigate this speculation by analyzing the population responses to natural movies and drifting grating in the same dataset.

**Figure S5:**
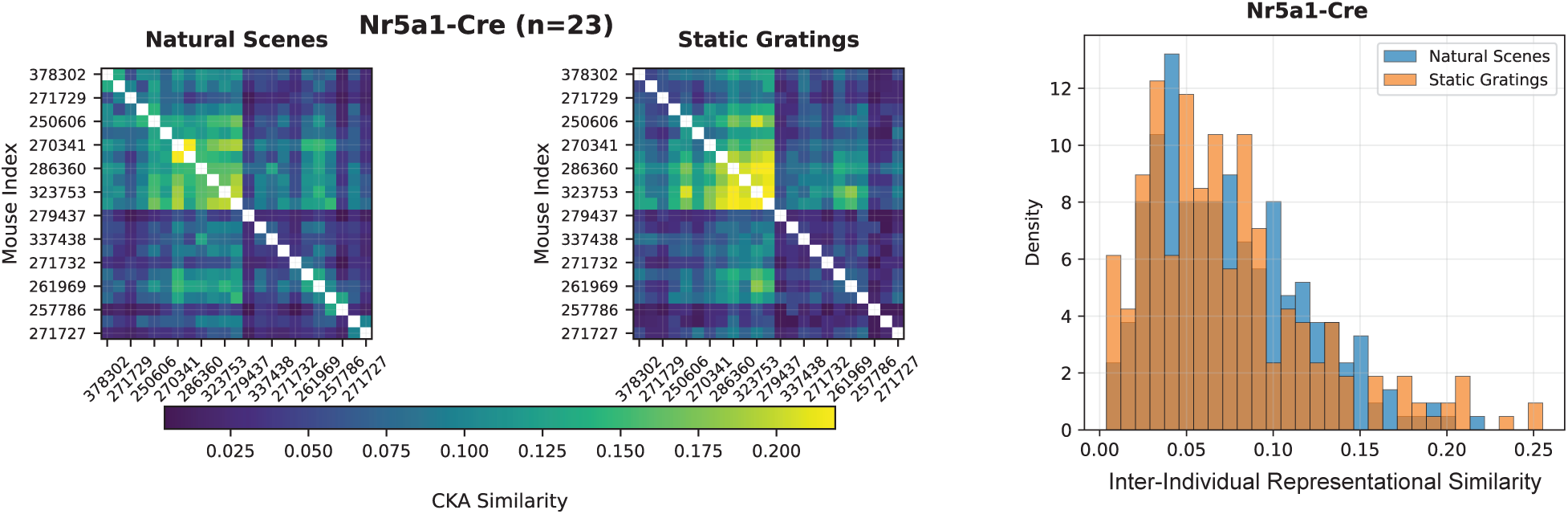
Nr5a1-expressing neurons show distinct coding patterns consistent with direction selectivity. Left panels show pairwise CKA similarity matrices between individual Nr5a1-expressing mice for natural scenes and static gratings (n=23 mice). Right panel shows the corresponding IIRS distribution histogram.

### B.4 Hierarchical clustering

Hierarchical clustering was performed on the condensed distance matrix (metric: linear CKA as in 2) using Ward’s linkage, yielding a dendrogram summarizing representational relationships across ¿100 region × layer × cell-type populations. Excitatory populations in V1 (red cluster) form a distinct, cohesive branch, suggesting that V1 excitatory circuits share a representational geometry distinct from higher-order areas. In contrast, inhibitory populations from V1 (SST, VIP, PV; blue, purple, orange) cluster with either inhibitory or excitatory populations from higher areas rather than with V1 excitatory populations, indicating that inhibitory circuitry exhibits conserved cross-area organization. This separation highlights that while excitatory V1 populations maintain locally specialized codes, inhibitory populations reflect more globally shared computational structure across the visual hierarchy. This pattern suggests that inhibitory circuits encode computations that generalize across cortical regions, while V1 excitatory neurons encode a locally specialized representational subspace for natural scenes.

**Figure S6:**
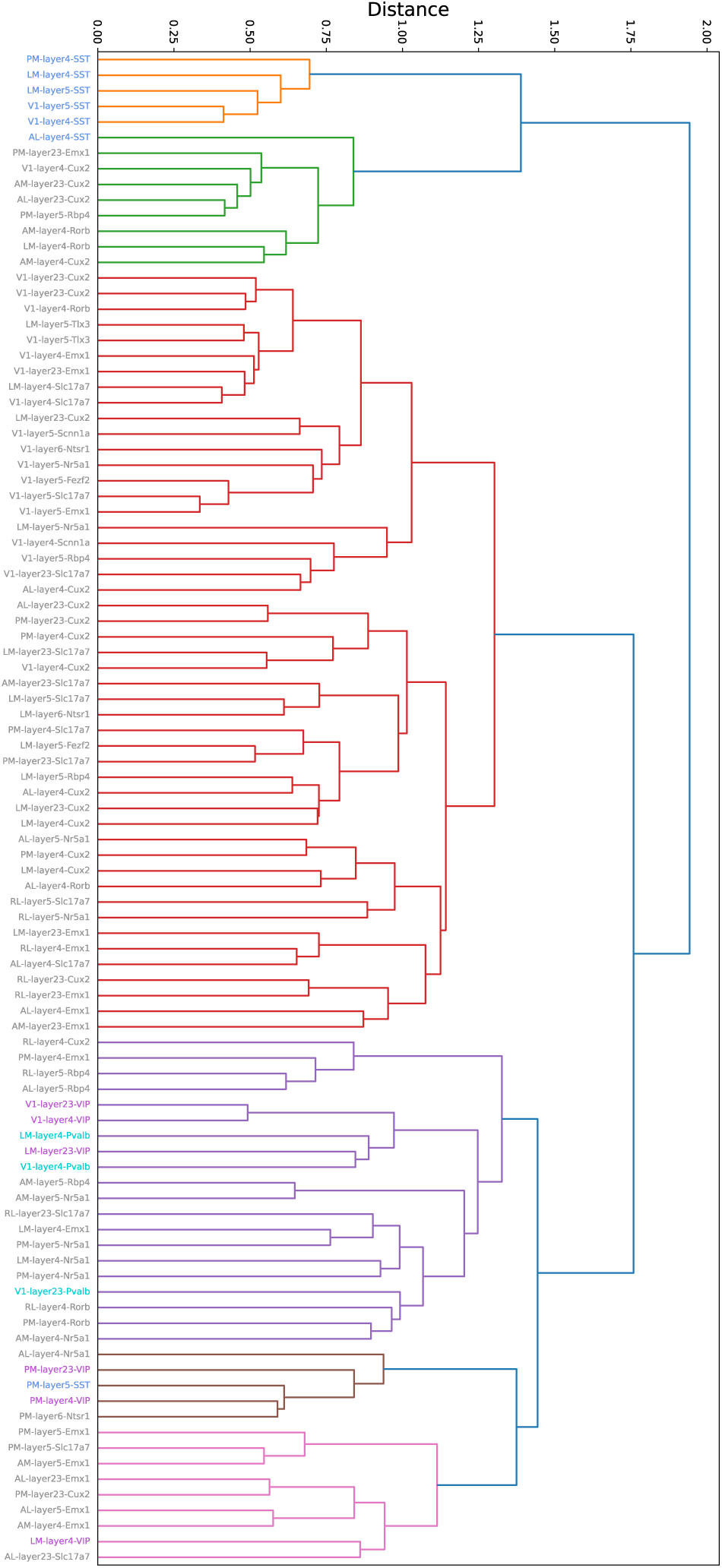
Hierarchical clustering for responses to natural scenes. Dendrogram showing the hierarchical organization of representational similarity (metric: linear CKA) across all combinations of genetically-defined cell type, cortical layer, and visual area. Each leaf corresponds to a population (e.g., V1-layer4-Cux2), computed from neurons with split-half reliability ¿ 0.4. Distances represent 1-CKA dissimilarity between population response matrices.

### B.5 Mapping cell-type-specific populations to task-optimized neural networks

While hierarchical clustering reveals the organizational structure of genetically-defined cell populations, it does not directly reveal what computational functions these clusters perform during visual processing. To bridge this gap between cellular identity and functional specialization, we systematically mapped neural population responses onto representations from deep networks trained on diverse visual tasks. We extracted representations of our natural stimulus set from deep neural networks trained on distinct visual tasks from the Taskonomy framework [32], including segmentation, texture edge detection, surface normal prediction, and keypoint detection. For each task-specific network, we computed representational similarity between network layer responses and cell-type population responses using Centered Kernel Alignment (CKA). This approach allows us to identify which computational features best align with different neural populations, effectively asking: “What visual computations does each cell type population specialize for?” Our analysis reveals a consistent functional organization that maps onto known anatomical pathways (Figure S7). Most remarkably, inhibitory neurons in V1 show computational profiles that are more similar to excitatory neurons in higher visual areas than to their local excitatory neighbors. For example, V1 inhibitory populations exhibit stronger alignment with surface normal and keypoint detection networks—computations typically associated with higher-order areas—while local V1 excitatory neurons show stronger segmentation similarity typical of early visual processing.

**Figure S7:**
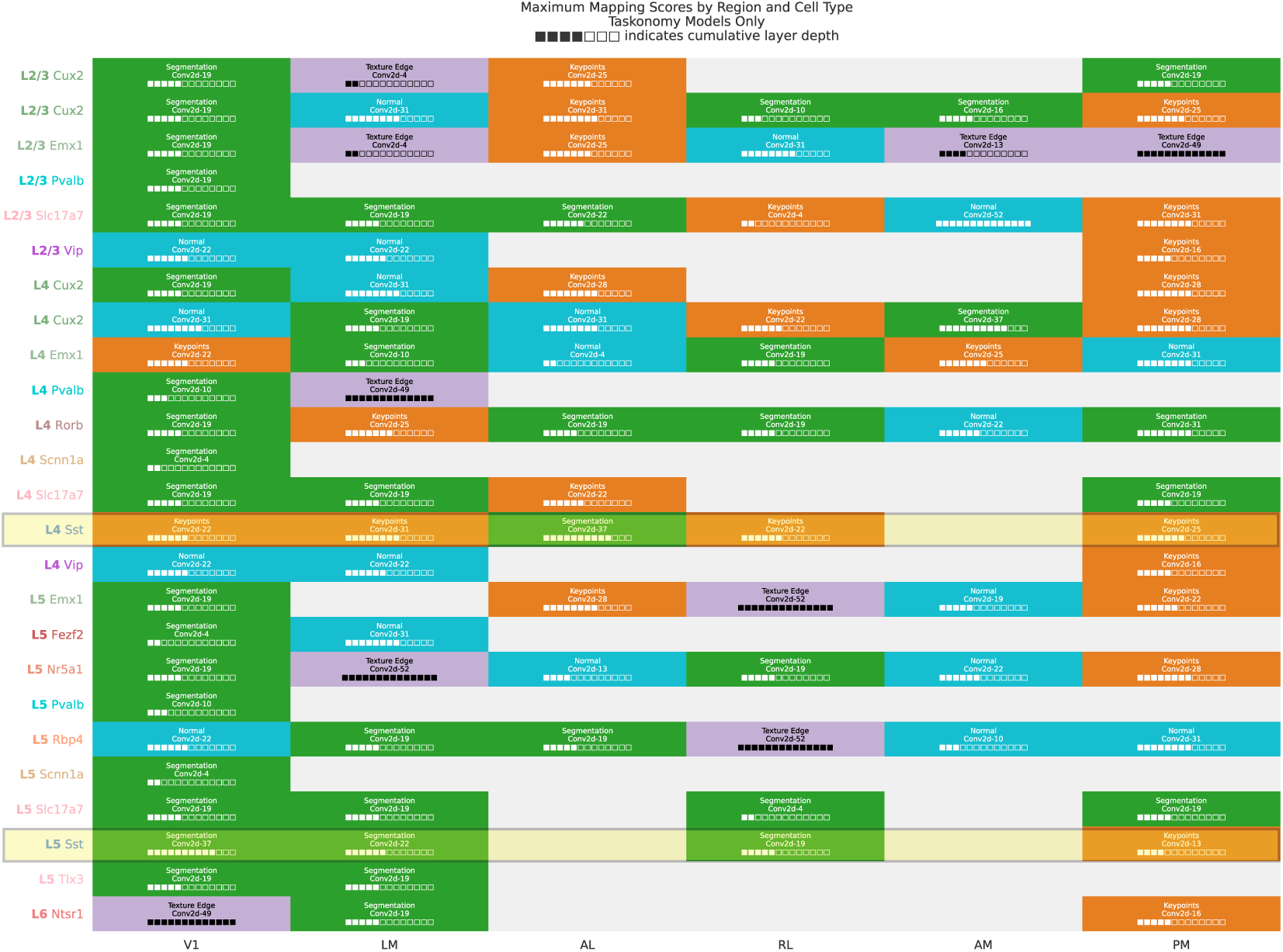
Functional Specialization of Genetically-Defined Cell Types Across Mouse Visual Areas. The figure shows the features with the highest representational similarity (CKA) between different Taskonomy model tasks (green: segmentation, purple: texture edge, blue: surface normals, and orange: key-points) and neural population responses across visual areas (V1, LM, AL, RL, AM, PM), layers (L2/3-L6), and genetic cell types (e.g., Cux2, Emx1). V1 and ventral area LM show stronger segmentation similarity, while dorsal pathway favors surface normals and keypoints detection. Sst cells were highlighted to show that we see a consistent pattern here as in UMAP, where Sst of L4-AL was different from the rest of Ssts of L4: keypoint detection vs segmentation.

